# SLC15A4 favors inflammasome function via mTORC1 signaling and autophagy restraint in dendritic cells

**DOI:** 10.1101/2022.03.15.484392

**Authors:** Cynthia López-Haber, Zachary Hutchins, Xianghui Ma, Kathryn E. Hamilton, Adriana R. Mantegazza

## Abstract

Phagocytosis is the first step in the assessment of foreign microbes or particles and enables activation of innate immune pathways such as the inflammasome. However, missing links between phagosomes and inflammasomes remain to be discovered. We show that in murine dendritic cells (DCs) the lysosomal histidine/peptide solute carrier transporter SLC15A4, associated with human inflammatory disorders, is recruited to phagosomes and is required for optimal inflammasome activity after infectious or sterile stimuli. Dextran sodium sulphate-treated SLC15A4-deficient mice exhibit decreased colon inflammation, reduced IL-1β production by intestinal DCs and increased autophagy. Similarly, SLC15A4-deficient DCs infected with *Salmonella* typhimurium show reduced caspase-1 cleavage and IL-1β production. This correlates with peripheral NLRC4 inflammasome assembly and increased autophagy. Overexpression of constitutively active mTORC1 rescues decreased IL-1β levels and caspase-1 cleavage, and restores perinuclear inflammasome positioning. Our findings suggest that SLC15A4 is a novel link that couples phagocytosis with inflammasome perinuclear assembly and inhibition of autophagy through phagosomal content sensing. Our data also reveal the previously unappreciated importance of mTORC1 signaling pathways to promote and sustain inflammasome activity.

## Introduction

Microorganisms or sterile particles captured by phagocytosis are first sensed by pattern-recognition receptors (PRRs) such as Toll-like receptors (TLRs), which recognize <u>p</u>athogen- or <u>d</u>anger-<u>a</u>ssociated <u>m</u>olecular <u>p</u>atterns (PAMPs or DAMPs) and are located at the cell surface and on phagosomes. Other PRRs localize to the cytosol and sense pathogen or phagocytosed cargo-derived products that escape phagosomes. In turn, activation of cytosolic PRRs – such as the <u>n</u>ucleotide binding domain leucine-rich repeat containing proteins (NLRs) – may lead to assembly of a multi-subunit complex known as the inflammasome and production of the highly inflammatory cytokine IL-1β, resulting in escalation of the immune response (Christgen & Kanneganti, 2020; Latz *et al*, 2013). Therefore, inflammasome activity must be tightly regulated to prevent chronic inflammation, which may cause excessive tissue damage and lead to disease. One negative regulator of inflammasomes is autophagy (Brady *et al*, 2018; Shi *et al*, 2012; Takahama *et al*, 2018; Zhong *et al*, 2016). We and others showed that inflammasome components such as the adaptor <u>a</u>poptosis-associated <u>s</u>peck-like protein containing a <u>c</u>aspase-recruitment domain (ASC) are targeted for autophagic sequestration (Mantegazza *et al*, 2017; Shi *et al*., 2012), leading to inflammasome silencing. However, the molecular mechanisms underlying the link between phagocytosis, inflammasome activity and autophagy are not completely understood (Moretti & Blander, 2014).

Phagosomes in dendritic cells (DCs) mature through a series of interactions with the endolysosomal system, acquiring increasing degradative capacity further promoted by phagosomal autonomous TLR signaling (Blander & Medzhitov, 2006; Hoffmann *et al*, 2012; Lopez-Haber *et al*, 2020). In addition to TLR recognition, phagolysosomal nutrient transporters may sense phagosomal degradation products and link nutrient sensing to the modulation of autophagy via the activation of <u>m</u>echanistic target <u>o</u>f rapamycin <u>c</u>omplex 1 (mTORC1) kinase signaling on phagolysosomes. mTORC1 senses lysosomal homeostasis and links cellular nutrient status to cell growth through its interactions with the Rag A-D GTPases and their regulators (Chantranupong *et al*, 2015; Perera & Zoncu, 2016). In nutrient-sufficient conditions, Rag complexes recruit mTORC1 to the lysosomal membrane for activation (Bar-Peled *et al*, 2012). Conversely, nutrient limitation inactivates Rag GTPases and consequently mTORC1. In turn, mTORC1 inactivation stimulates autophagy by relieving inhibition of the master lysosomal/ autophagy gene transcriptional activators (Martina *et al*, 2012; Raben & Puertollano, 2016; Sardiello *et al*, 2009; Settembre *et al*, 2012). Importantly, dysregulation of nutrient sensing may lead to disease, as shown by the association of many solute transporters with human metabolic and inflammatory disorders such as gout, <u>s</u>ystemic lupus <u>e</u>rythematosus (SLE) and inflammatory <u>b</u>owel <u>d</u>isease (IBD) (Lin *et al*, 2015; Zhang *et al*, 2019).

Among phagolysosomal <u>s</u>olute <u>c</u>arrier (SLC) transporters is the histidine/peptide transporter SLC15A4. In plasmacytoid DCs (pDCs) and B cells – in which this transporter has been extensively studied – SLC15A4 is required for optimal endosomal TLR7/9 signaling and subsequent type I interferon and antibody production (Blasius *et al*, 2010; Kobayashi *et al*, 2021a; Sasai *et al*, 2010). In line with this, SLC15A4 is associated with SLE and IBD in genome-wide association studies (Baccala *et al*, 2013; Heinz *et al*, 2020; Kobayashi *et al*., 2021a; Kobayashi *et al*, 2014; Sasawatari *et al*, 2011). Moreover, increased SLC15A4 mRNA levels were detected in colon samples from a cohort of IBD patients (Lee *et al*, 2009). Interestingly, in conventional DCs – in which SLC15A4 has been much less studied – it allows bacterially derived peptidoglycan egress from endosomes to promote pro-inflammatory signaling through the cytosolic sensors <u>n</u>ucleotide-binding <u>o</u>ligomerization <u>d</u>omain 1/2 (NOD1/2) (Nakamura *et al*, 2014), also associated with IBD (Hugot *et al*, 2001; Sasawatari *et al*., 2011). However, the potential role of SLC15A4 in inflammasome activity has not been explored. Of note, SLC15A4 bears a di-leucine motif recognized by the endolysosomal adaptor protein-3 (AP-3), which promotes cargo sorting to lysosomes and lysosome-related organelles (Blasius *et al*., 2010). Our previous studies demonstrated that AP-3 is required for optimal phagosomal TLR signaling in DCs and promotes inflammasome priming (Lopez-Haber *et al*., 2020; Mantegazza *et al*, 2012; Mantegazza *et al*., 2017; Mantegazza *et al*, 2014). In addition, AP-3 indirectly regulates NLRC4 inflammasome positioning in the perinuclear region to prevent autophagic sequestration and limits autophagy to sustain inflammation and control bacterial infection (Mantegazza *et al*., 2017). We hypothesized that the phenotype associated with AP-3 deficiency might be partly attributed to its putative cargo SLC15A4. We now show that SLC15A4 is a novel link between phagocytosis and inflammasome activation by coupling phagosomal content sensing to mTORC1 signaling. We demonstrate that SLC15A4 promotes inflammasome activity after *Salmonella enterica* serovar Typhimurium (STm) infection or sterile particle stimulation by restraining autophagy both *in vitro* and in an *in vivo* model of dextran sodium sulphate (DSS)-induced colitis. Additionally, our data unravel a heretofore unappreciated role for SLC15A4 and mTORC1 in sustaining NLRC4 inflammasome activity by ensuring proper complex assembly at the perinuclear region and limiting autophagy.

## Results

### SLC15A4 promotes inflammasome activity by inhibiting autophagy after phagocytosis

We previously showed that AP-3 is required for optimal phagosomal maturation and formation of phagosomal tubules upon phagocytosis in DCs, which is dependent on the recruitment of lipid kinase phosphatidylinositol-4-kinase-2alpha to phagosomes (Lopez-Haber *et al*., 2020; Mantegazza *et al*., 2014). We also showed that AP-3 indirectly sustains NLRP3 and NLRC4 inflammasome activity induced by phagosomal stimuli (Mantegazza *et al*., 2017). We propose that this function is likely due to AP-3-dependent recruitment of other unknown cargo to phagosomes. Given that SLC15A4 may bind AP-3 (Blasius *et al*., 2010), we first investigated the recruitment of SLC15A4 to DC phagosomes after phagocytosis of polystyrene beads coated with bacterial lipopolysaccharide (LPS) and Texas red (TxR)-conjugated ovalbumin (OVA), as well as STm. We observed that SLC15A4-GFP is indeed recruited to phagosomes and phagosomal tubules after LPS/OVA bead phagocytosis in wild-type (WT) bone marrow-derived DCs (BMDCs) and the DC line DC2.4 (**Fig. 1A**, **Fig. EV1** and **Movies EV1 and EV2**), as well as to STm-containing phagosomes (**Fig. 1B**).

**Figure 1.**
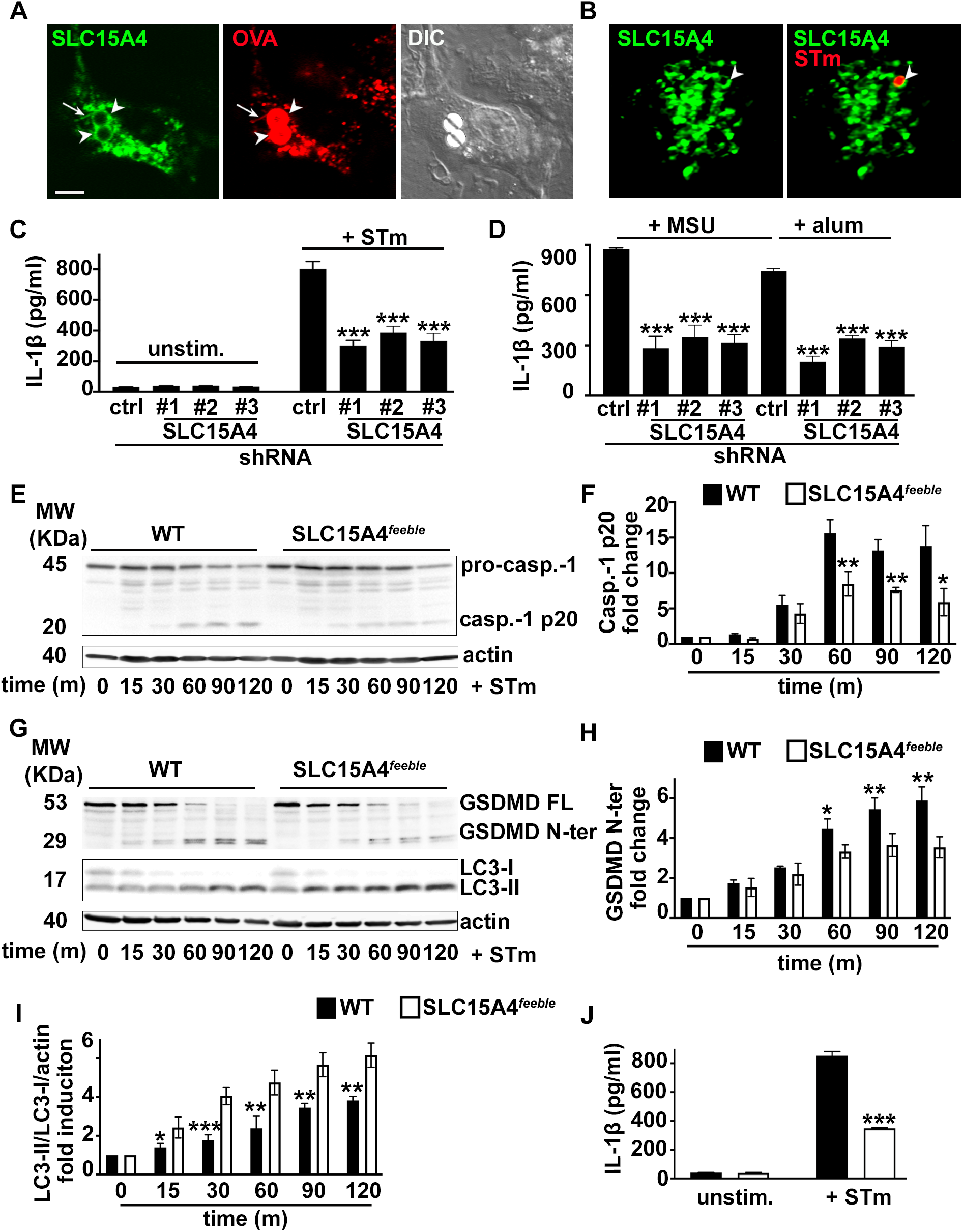
SLC15A4 promotes inflammasome activity by inhibiting autophagy. WT BMDCs transduced with retrovirus encoding SLC15A4-GFP (**A, B**), or lentiviruses encoding non-target (ctrl) or any of three SLC15A4-specific shRNAs (**C, D**) or WT or SLC15A4*^feeble^* BMDCs (**E-J**) were treated with LPS/OVA-TxR polystyrene beads (**A**) or STm-mcherry (**B**), unstimulated or infected with flagellin-expressing STm (**C, E-J**) or primed for 3 h with soluble LPS and stimulated with alum or MSU (**D**). (**A, B)**. Cells were analyzed by live cell imaging 2 h after the pulse (**A**) or 1 h after infection (**B**). Representative images. Differential interference contrast (DIC) image shows cell shape and outline. Arrowheads, phagosomes; arrows, phagosomal tubules. Scale bar, 6 μm. (**C-J)**. Cell supernatants collected 2 h (**C, J**) or 4 h (**D**) after treatment were assayed for IL-1β by ELISA. (**C, D, J**). Representative plots of 3 independent experiments. (**E-I**). Cell pellets collected at the indicated time points after STm infection were lysed, fractionated by SDS-PAGE and immunoblotted for caspase-1 (**E, F**), GSDMD or LC-3 (**G-I**). (**E, G**). Representative immunoblots showing pro-caspase-1 (pro-casp.-1) and cleaved p20 (casp.-1 p20) bands (**E**) or GSDMD full length (GSDMD FL), cleaved GSDMD N-terminal fragment (GSDMD N-ter), LC3-II and LC3-I bands (**G**). (**F, H, I**) Quantification of band intensities for caspase-1 p20 normalized to pro-caspase-1 and actin (**E**), GSDMD N-ter normalized to GSDMD FL and actin (**G**) and LC3-II normalized to LC3-I and actin (**H**) from three independent experiments are shown as fold change (**F, H**) or fold induction (**I**) relative to time 0. Data represent mean ± SD. * p<0.05; **p<0.01; ***p<0.001. See also **Figs. EV1 and EV2**.

We then investigated whether SLC15A4 could regulate inflammasome activity after phagocytosis. To test this, we knocked-down SLC15A4 in BMDCs using three different shRNA sequences. Knock-down efficiency was between 50 and 80%, as assessed by quantitative real-time PCR (**Fig. EV2A**). SLC15A4 knockdown (KD) did not affect DC differentiation compared to shRNA non-target control-treated DCs (**Fig. EV2C**). We then treated BMDCs with flagellin-expressing STm – to induce NLRC4 activation (Mantegazza *et al*., 2017) – and measured IL-1β secretion as a read-out for inflammasome activity by ELISA. IL-1β secretion was reduced by 50% in SLC15A4 KD DCs relative to shRNA control-treated DCs (**Fig. 1C**). Similar results were observed when DCs were primed with LPS and subsequently stimulated with alum or monosodium urate crystals (MSU) to activate the NLRP3 inflammasome (**Fig. 1D**). Conversely, secretion of IL-6 after LPS treatment was not affected by SLC15A4 KD, suggesting that SLC15A4 does not play a role in the priming step of the inflammasome pathway (**Fig. EV2B**). This is in agreement with previous observations indicating that SLC15A4 does not affect TLR signaling in conventional DCs(Blasius *et al*., 2010). To confirm our observations on SLC15A4 KD DCs, we differentiated DCs from WT and SLC15A4*^feeble^* mice. The *feeble* mutation results in abnormal splicing of the *Slc15a4* gene resulting in the transcription of two aberrant products and the absence of functional protein expression(Blasius *et al*., 2010). SLC15A4*^feeble^* BMDCs differentiated and maturated similarly in response to LPS compared to BMDCs (**Fig. EV2D, E**). In agreement with our observations in SLC15A4 KD DCs, after STm stimulation, IL-1β secretion was reduced by more than 50% in SLC15A4*^feeble^* DCs compared to WT DCs (**Fig. 1J**). Consistent with the decreased IL-1β secretion, pro-caspase 1 and its substrate gasdermin-D (GSDMD) cleavage were also reduced in SLC15A4*^feeble^* DCs between 60 and 120 minutes after STm stimulation (**Fig. 1E-H**). These observations suggest that SLC15A4 is required for optimal inflammasome activity.

We previously showed that STm infection induces autophagy in BMDCs and that the absence of AP-3 increases autophagy induction (Mantegazza *et al*., 2017). However, the mechanism by which AP-3 modulates autophagy remains unknown. Considering the link between SLC15A4 and mTORC1 – a known autophagy regulator (Martina *et al*., 2012; Settembre *et al*., 2012)– in other cell types (Kobayashi *et al*., 2014), we hypothesized that SLC15A4 may be required for optimal inflammasome activity in DCs by limiting autophagy. We detected induction of the lipidated form of the microtubule-associated protein 1 light <u>c</u>hain <u>3</u> α (LC3-II) relative to the unlipidated LC3-I (indicative of autophagy induction (Klionsky *et al*, 2021)), in cell lysates overtime after STm stimulation in WT DCs (**Fig. 1G, I**). Importantly, LC3-II induction was significantly increased in SLC15A4*^feeble^* DCs between 15 and 120 minutes after STm stimulation compared to WT DCs (**Fig. 1G, I**). These observations suggest that SLC15A4 promotes inflammasome activity by inhihiting autophagy after STm stimulation.

### SLC15A4*^feeble^* mice show faster recovery from acute dextran sodium sulphate-induced colitis in mice

SLC15A4 expression was shown to be pro-colitogenic in a mouse model of mild chronic DSS-induced colitis due to its role in promoting TLR9 and NOD signaling (Sasawatari *et al*., 2011). However, the role played by SLC15A4 in inflammasome activity and the production of IL-1β in DCs has not been investigated. To test whether SLC15A4 regulates inflammasome activity *in vivo*, we employed a model of acute DSS-induced colitis. This model is proposed to cause intestinal injury associated with NLRP3 inflammasome activation and IL-1β production (Bauer *et al*, 2010). WT and SLC15A4*^feeble^* mice received either water (naïve mice) or 2.5% DSS in drinking water for 5 days, followed by water for 10 more days to allow recovery. Weight loss and stool appearance, consistency and presence of blood were monitored daily after DSS administration. WT and SLC15A4*^feeble^* mice showed a similar trend of body weight loss during the first 8 days after the start of DSS administration – with an average of 15% and 13% body weight loss on day 7, respectively. However, after this time point, SLC15A4*^feeble^* mice recovered body weight significantly more efficiently than WT mice. Remarkably, at day 16 (endpoint), SLC15A4*^feeble^* mice completely recovered their initial body weight, while WT mice body weight remained significantly lower – 0% and 10% of body weight loss, respectively (**Fig. 2A**). Even though the percent of body weight loss was similar at day 8, the blood stool index was significantly different between WT and SLC15A4*^feeble^* mice. Whereas most of WT mice exhibited very soft stool consistency and visible traces of stool blood (score 2), most of SLC15A4*^feeble^* mice showed soft but formed stool consistency and positive hemoccult tests without visible traces of blood (score 1) (**Fig. 2B**, ***left*)**. At day 16, consistent with the faster recovery observed in body weight in SLC15A4*^feeble^* mice, most of these mice exhibited normal stool appearance and negative hemoccult tests (score 0), whereas most of WT mice remained showing higher blood stool index scores at the same time point (**Fig. 2B**, ***right***).

**Figure 2.**
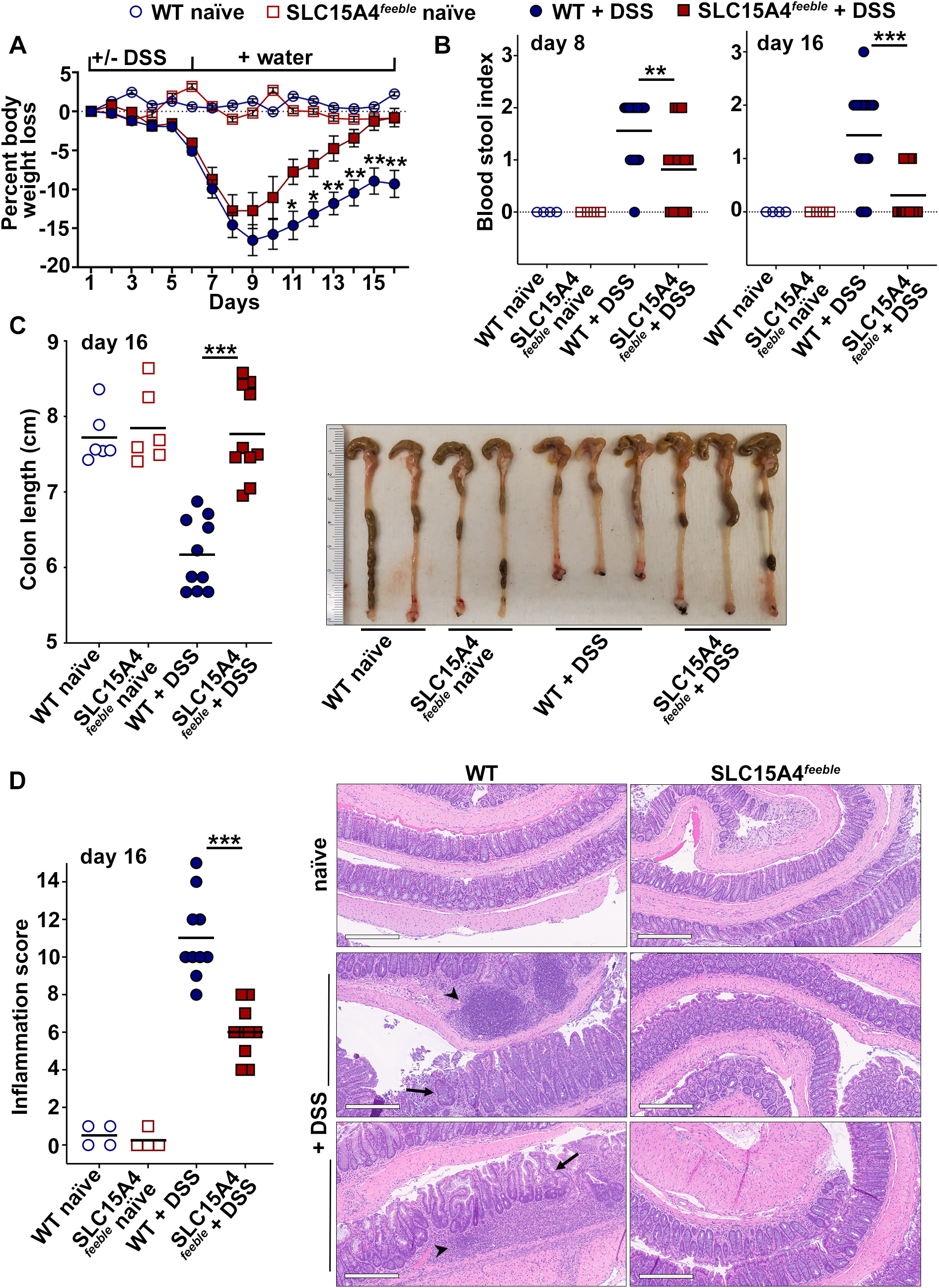
SLC15A4*^feeble^* mice show faster recovery from acute dextran sodium sulphate-induced colitis in mice. WT and SLC15A4*^feeble^* mice were given 2.5% DSS in drinking water for 5 days followed by normal drinking water for 10 more days or water only (naïve). **A**. Mice were weighed daily. Percent of body weight loss overtime is represented relative to day 1. **B**. Stool consistency and presence of blood were assessed. Blood stool index at days 8 (*left panel*) and 16 (*right panel*) was determined using the following score: 0, normal feces, negative hemoccult; 1, soft but formed feces, positive hemoccult; 2, very soft feces, visible traces of stool blood; 3, diarrhea, rectal bleeding. **C**. *Left panel*. Colon length was measured at day 16. *Right panel*. Representative image of 3 independent experiments. **D**. Colon swiss roll samples were formalin-fixed, paraffin embedded, stained and scored blinded by expert pathologist. *Left panel*. Inflammation score at day 16. *Right panels*. Representative hematoxylin and eosin stained, formalin-fixed paraffin embedded samples. Note disrupted crypt architecture (*arrows*) and inflammatory infiltrate (*arrowheads*) in DSS-treated WT samples. Data represent mean ± SD. * p<0.05; **p<0.01; ***p<0.001; Mann-Whitney non-parametric statistical test.

Decreased colon length has been associated with increased inflammation (Chassaing *et al*, 2014). In agreement with this observation and with our previous results at day 16, while most of DSS-treated SLC15A4*^feeble^* mice showed similar colon length compared to water-treated mice (mean= 7.8 cm), all WT mice showed significantly reduced colon length compared to water-treated mice and DSS-treated SLC15A4*^feeble^* mice (mean= 6.2 cm) (**Fig. 2C**), suggesting that colon inflammation is reduced in the absence of SLC15A4. Consistent with these observations, histopathological evaluation of swiss roll samples prepared from mouse colon at day 16, revealed significant differences between WT and SLC15A4*^feeble^* mice (**Fig. 2D**). WT mouse samples displayed in most cases moderate to severe inflammation characterized by focally extensive ulceration of the mucosa with segmental involvement of the deep lamina propria and multifocal extension into the submucosa, while SLC15A4*^feeble^* mouse samples showed mostly mild to moderate inflammation with modest infiltrate of mononuclear cells (**Fig. 2D, *right***). Accordingly, the inflammation score index – comprising mucosal/crypt loss, crypt inflammation and inflammatory infiltrate – was remarkably higher in WT samples (mean= 11) compared to SLC15A4*^feeble^* samples (mean= 6) (**Fig. 2D, *left***). These results show that the absence of SLC15A4 promotes faster recovery from acute DSS-induced colitis.

Given our observations that SLC15A4 favors IL-1β production in BMDCs by restraining autophagy, we assessed the contribution of intestinal DCs to the DSS-induced process. At day 16, DCs from WT and SLC15A4*^feeble^* mice were isolated from colon. The percent of CD103+/CD11c+ DCs was similar between WT and SLC15A4*^feeble^* mice (**Fig. EV2F**). Isolated DCs were then cultured overnight without further stimulation. Remarkably, IL-1β secretion was significantly higher in the cell culture supernatants from WT DCs relative to SLC15A4*^feeble^* DCs as measured by ELISA (**Fig. 3A**). In agreement with our *in vitro* results, cell lysates from isolated intestinal DCs showed increased LC3-II induction in SLC15A4*^feeble^* DCs compared to WT DCs (**Fig. 3B**), suggesting that increased autophagy is limiting IL-1β production in colonic DCs *in vivo*.

**Figure 3.**
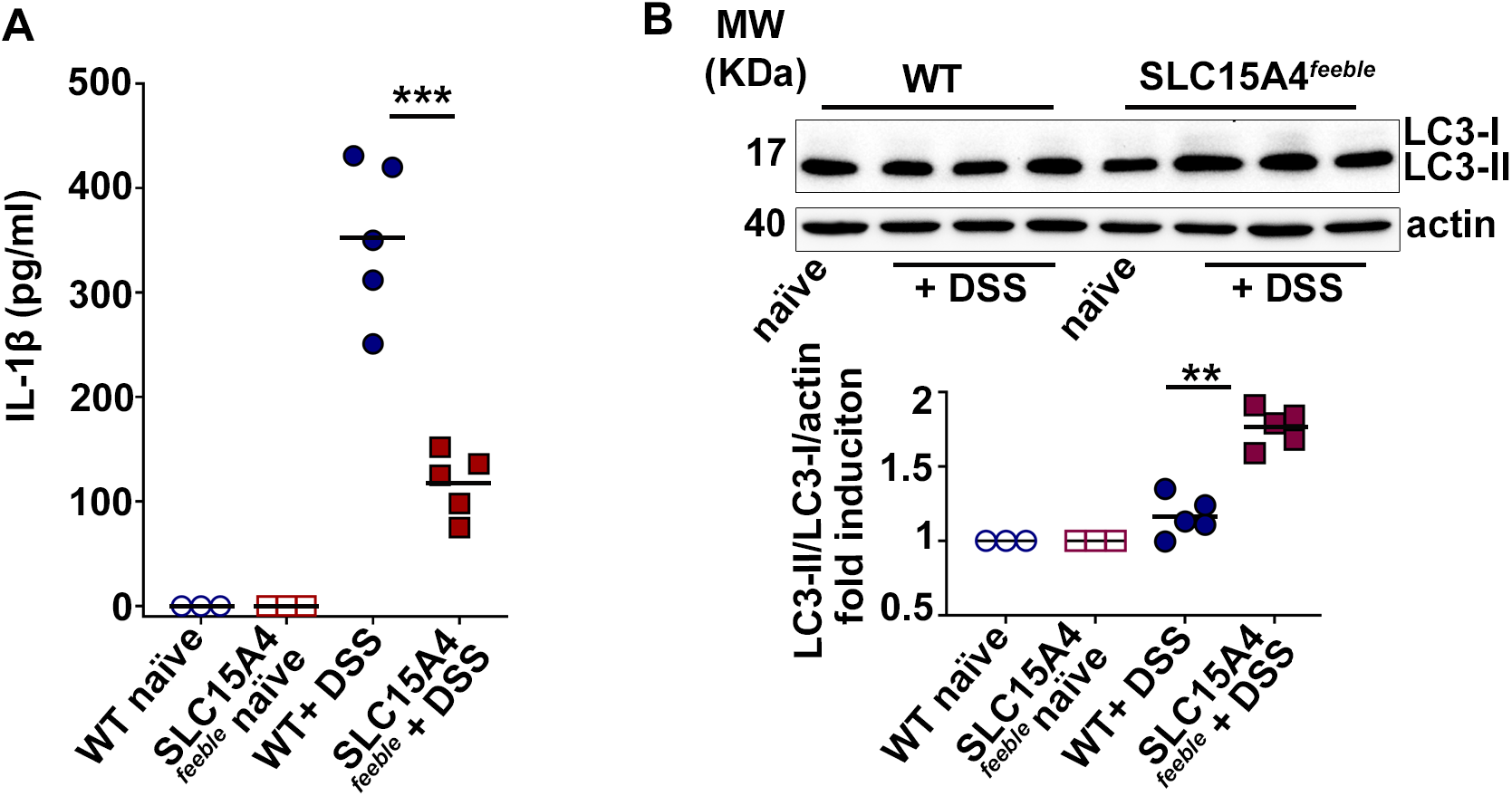
SLC15A4*^feeble^* intestinal CD11c+ DCs show reduced IL-1β secretion and increased autophagy. Intestinal CD11c+ DCs were isolated from colon of WT and SLC15A4*^feeble^* mice at day 16 and cultured overnight. **A**. IL-1β was measured from culture supernatants by ELISA. **B**. Cell pellets were immunoblotted for LC3 and actin. *Upper panel*. Representative immunoblot. *Lower panel*. Quantification of band intensities for LC3-II normalized to LC3-I and actin are shown as fold induction relative to samples from untreated mice. Note that LC3-I signal from *ex vivo* samples is almost undetectable. Data represent mean ± SD. ***p<0.001; **p<0.01. Mann-Whitney non-parametric statistical test. See also **Fig. EV2F**.

### SLC15A4 promotes inflammasome perinuclear positioning away from autophagic membranes

We and others showed that autophagy negatively regulates inflammasome activity and that the adaptor ASC aggregates or “specks” formed after inflammasome assembly are sequestered by autophagic membranes (Mantegazza *et al*., 2017; Shi *et al*., 2012). To assess inflammasome assembly and ASC speck sequestration, we transduced WT and SLC15A4*^feeble^* DCs with the retroviral constructs ASC-GFP and mcherry-LC3 or probed endogenous ASC specks together with the autophagic adaptor p62/SQTM1 (p62), which links ubiquitylated substrates to LC3 (Clausen *et al*, 2010). ASC speck formation was monitored by fluorescence microscopy and flow cytometry. Neither the percentage of ASC-GFP speck-positive DCs nor the kinetics of ASC-GFP speck formation after STm infection differed appreciably between WT and SLC15A4*^feeble^* DCs as assessed by flow cytometry by analyzing GFP-width and GFP-height parameters as previously described (Hoss *et al*, 2018; Sester *et al*, 2015) (**Fig. EV3**).

On the contrary, ASC-GFP specks in SLC15A4*^feeble^* DCs were increasingly surrounded by LC3-II puncta (within a radius of 1 μm), compared to WT DCs as observed by fluorescence microscopy 30 minutes after STm stimulation (**Fig. 4A, B**). Similarly, endogenous ASC specks in SLC15A4*^feeble^* DCs were also significantly more surrounded by p62 puncta at the same time point (**Fig. 4C**). Additionally, ASC speck formation was visualized mostly in the perinuclear region (within a radius of 3 μm), in WT DCs (70 ± 10% of cells), while intriguingly, in most of SLC15A4*^feeble^* DCs it was detected away from the nucleus (26 ± 9% of cells in perinuclear region; **Fig.4A, C, D**). This behavior resembles our previous observations in AP-3-deficient DCs, in which ASC specks appear away from the nucleus and surrounded by autophagic membranes(Mantegazza *et al*., 2017).

**Figure 4.**
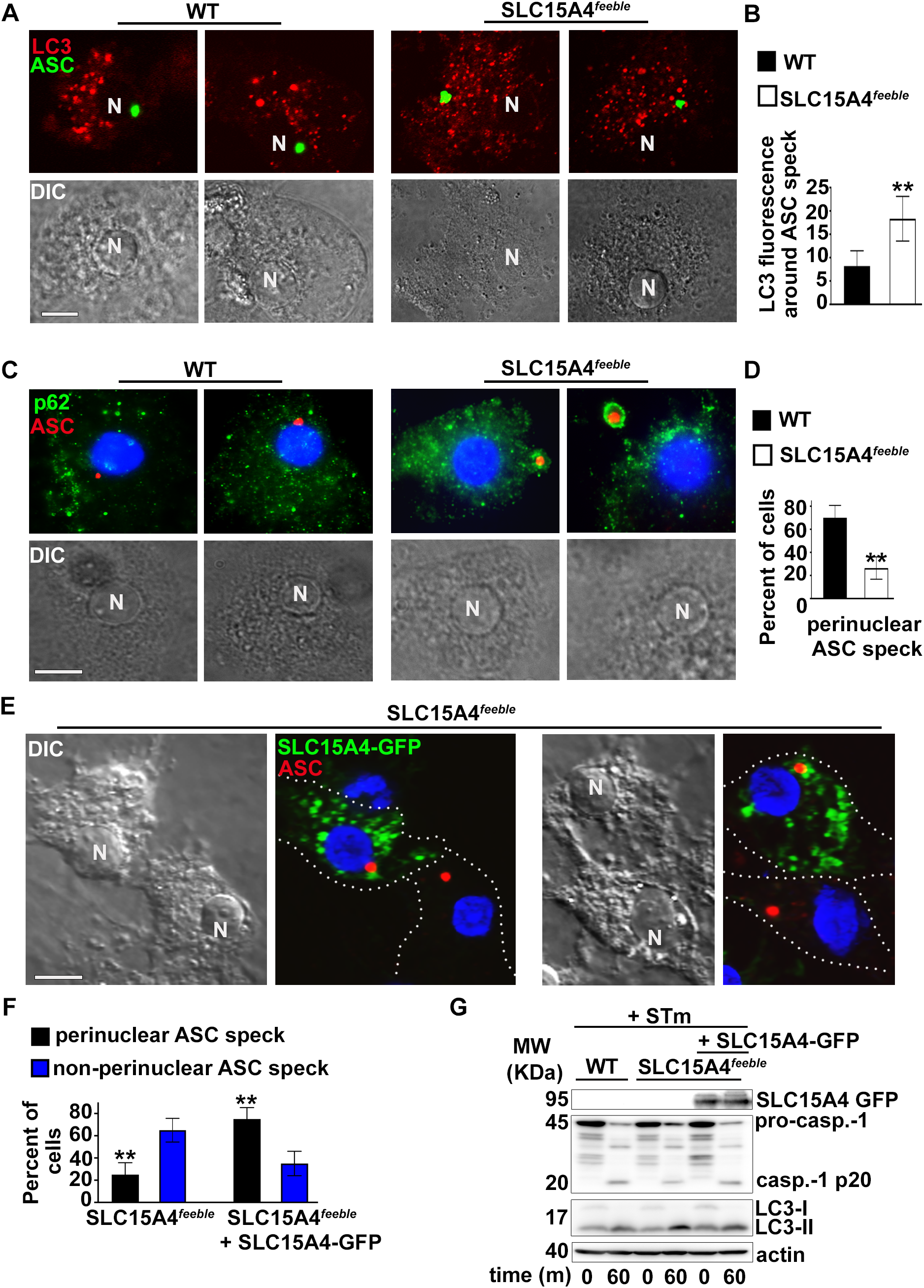
SLC15A4 promotes inflammasome perinuclear positioning away from autophagic membranes. WT or SLC15A4*^feeble^* BMDCs expressing ASC-GFP and mCherry-LC3 (**A, B, D**) or non-transduced (**C**) or SLC15A4*^feeble^* BMDCs transduced with SLC15A4-GFP (**E**, **F, G**) were infected with flagellin-expressing STm and fixed (**A-F**) or lysed (**G**) 1 h after infection. **A**. Cells were analyzed by fluorescence microscopy. Representative images showing ASC speck (green) relative to mCherry-LC3 (red) in two infected WT and SLC15A4*^feeble^* DCs each. **B.** Quantification of LC3 fluorescence per unit area in a radius of 1 μm surrounding the ASC speck in 20 cells per cell type in each of 3 independent experiments. **C.** Cells were stained for endogenous ASC and p62, labeled with DAPI and analyzed by fluorescence microscopy. Representative images showing ASC speck (red) relative to p62 (green) in two infected WT and SLC15A4*^feeble^* DCs each. Note p62 staining surrounding ASC specks in SLC15A4*^feeble^* DCs. **D.** Quantification of perinuclear (within a radius of 3 μm from the nucleus) ASC specks in 30 cells per cell type in each of 3 independent experiments. (**E**, **F**). Cells were stained for endogenous ASC, labeled with DAPI and analyzed by fluorescence microscopy. Representative images showing ASC speck (red) and SLC15A4-GFP (green) in SLC15A4*^feeble^* DCs together with non-transduced SLC15A4*^feeble^* DCs. Note the perinuclear positioning of ASC specks in the transduced cells (**E**). Quantification of perinuclear (within a radius of 3 μm from the nucleus) and non-perinuclear ASC specks in 30 cells per cell type in each of 3 independent experiments (**F**). **G**. Representative immunoblots showing pro-caspase-1 (pro-casp.-1) and cleaved p20 (casp.-1 p20), LC3-II and LC3-I, and actin bands. N, nucleus. Dotted white lines, cell outlines. Corresponding DIC images show nuclear position. Scale bar, 8 μm. Data represent mean ± SD. **p<0.01. See also **Fig. EV3**.

ASC speck positioning in SLC15A4*^feeble^* DCs shifted from peripheral to perinuclear in cells transduced with human SLC15A4-GFP (**Fig. 4E, F**). This correlated with increased inflammasome activity as evidenced by augmented caspase-1 cleavage in SLC15A4*^feeble^* DCs expressing human SLC15A4-GFP compared to non-transduced counterparts (**Fig. 4G**).

Altogether, these observations suggest that the kinetics of inflammasome formation is not affected by SLC15A4. However, the site of inflammasome assembly, inflammasome activity and its targeting by autophagy is regulated by SLC15A4.

### Constitutive mTORC1 signaling restores inflammasome activity and positioning in SLC15A4*^feeble^* DCs

SLC15A4 was shown to regulate mTORC1 signaling in B cells and mast cells (Kobayashi *et al*., 2014; Kobayashi *et al*, 2017). Given that mTORC1 activation inhibits autophagy (Martina *et al*., 2012; Settembre *et al*., 2012), we investigated whether mTORC1 signaling was dysregulated in SLC15A4*^feeble^* DCs in our model of *in vitro* STm infection, in which SLC15A4 restrains autophagy (**Fig. 1G, I**). WT and SLC15A4*^feeble^* DCs non-transduced or transduced with the control construct methionine aminopeptidase 2 (metap2 (Gu *et al*, 2017) or GTP-bound RagB^Q99L^ – which lacks GTPase activity and renders mTORC1 constitutively active (Bar-Peled *et al*., 2012)– were infected with STm for 1h and assayed for mTORC1 activity. We observed ∼50% reduced phosphorylation of mTOR on Ser 2481 – the autophosphorylation site associated with mTORC1 catalytic activity(Kobayashi *et al*., 2014) – together with a similar impaired phosphorylation of mTORC1 downstream substrates p70S6 kinase and S6 ribosomal protein in infected SLC15A4*^feeble^* DCs compared to their WT counterparts (**Fig. 5A, B**; non-transduced and control lanes). Reduced phosphorylation in SLC15A4*^feeble^* DCs was rescued when DCs were transduced with GTP-bound RagB^Q99L^ (**Fig. 5A, B**). Similar results were obtained after cell starvation in Earle’s balanced salt solution (EBSS), which lacks essential aminoacids (Sharifi *et al*, 2015) (**Fig. EV4A**). Notably, reduced IL-1β production together with decreased caspase-1 and GSDMD cleavage in STm infected SLC15A4*^feeble^* DCs were also rescued by overexpression of RagB^Q99L^ (**Fig. 5C-E**). Conversely, inhibition of mTOR signaling in WT BMDCs using torin, reduced IL-1β production to the levels detected in SLC15A4*^feeble^* DCs (**Fig. EV4B**). These observations suggest that SLC15A4 positively regulates mTORC1 signaling in DCs, keeping autophagy at bay to promote inflammasome activity after STm infection.

**Figure 5.**
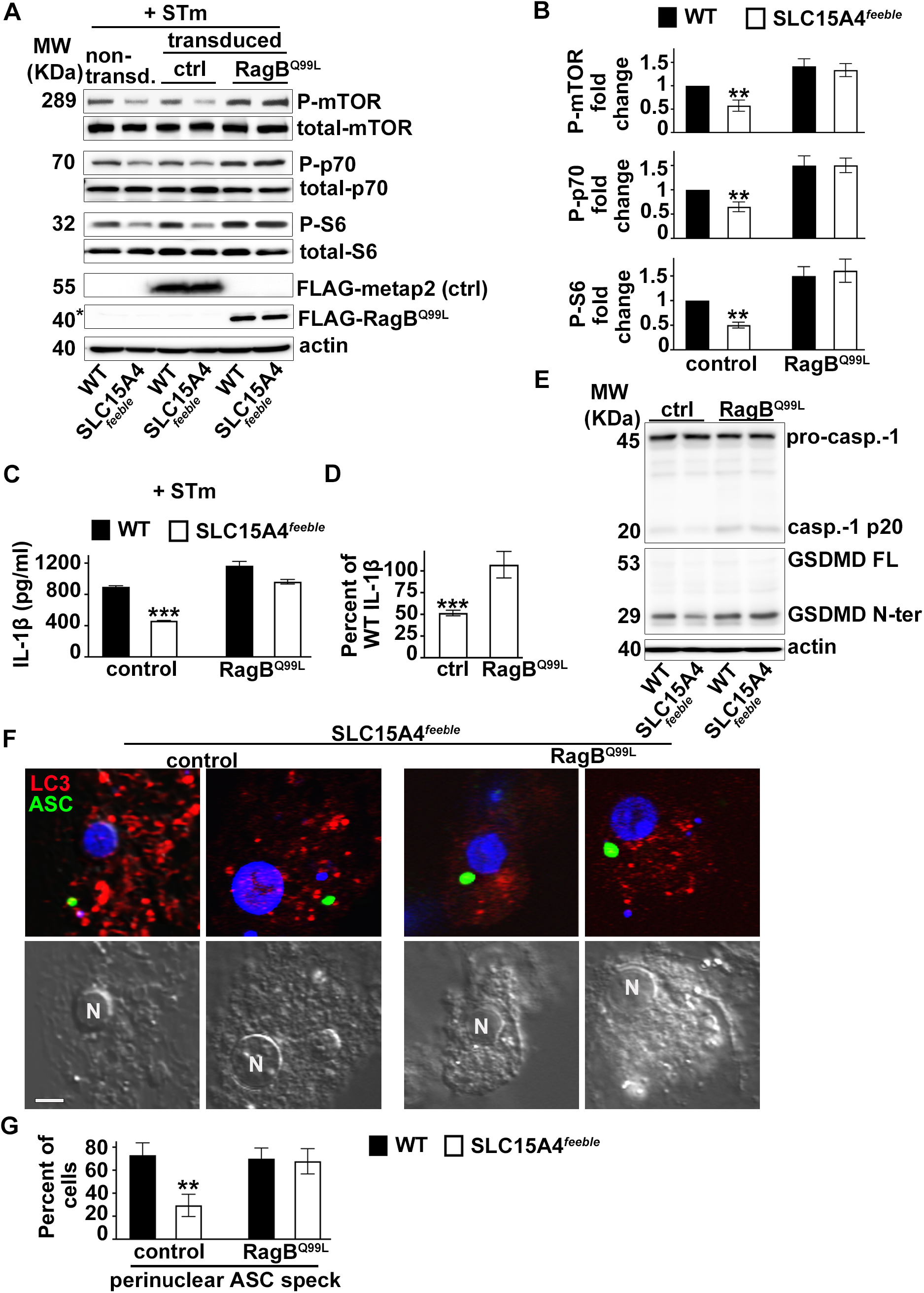
Constitutive mTORC1 signaling restores inflammasome activity and positioning in SLC15A4*^feeble^* DCs. WT or SLC15A4*^feeble^* BMDCs transduced with constitutively active RagB (RagB^Q99L^) or metap2 (control) (**A-G**) and mcherry-LC3 (**F, G**) were infected with flagellin-expressing STm. (**A**, **B**). Cell pellets collected 1 h after STm infection were lysed, fractionated by SDS-PAGE and immunoblotted for phospho (P) and total mTOR, p70 kinase and S6, FLAG and actin. Representative immunoblots. Non-specific band right above FLAG-RagB^Q99L^ is indicated with an asterisk (**A**). Quantification of band intensities for P-mTOR normalized to total mTOR, P-p70 normalized to total p70 and P-S6 normalized to S6 (**B**). (**C, D**). Cell supernatants collected 1 h after infection were assayed for IL-1β by ELISA. **C**. Representative plot of 3 independent experiments. **D**. IL-1β values of transduced SLC15A4*^feeble^* BMDCs from 3 independent experiments are shown as percent of WT DC values to represent rescue of the defective phenotype. **E.** Cell pellets collected 1 h after STm infection were probed for caspase-1 and GSDMD. Representative immunoblots showing pro-caspase-1 (pro-casp.-1) and cleaved p20 (casp.-1 p20), GSDMD full length (GSDMD FL) and cleaved GSDMD N-terminal fragment (GSDMD N-ter) bands. (**F, G**). Cells were stained for endogenous ASC, labeled with DAPI and analyzed by fluorescence microscopy. Representative images showing ASC speck (green) and LC3 (red) in SLC15A4*^feeble^* DCs (**F**). Quantification of perinuclear (within a radius of 3 μm from the nucleus) in 30 cells per cell type in each of 3 independent experiments (**G**). N, nucleus. Corresponding DIC images show nuclear position. Scale bar, 8 μm. Data represent mean ± SD. **p<0.01; ***p<0.001. See also **Fig. EV4**.

Remarkably, overexpression of RagB^Q99L^, but not metap2 (control), in SLC15A4*^feeble^* DCs induced ASC speck formation in the perinuclear region after NLRC4 stimulation (**Fig. 5F, G**), a location that appears to be protected from autophagic membranes (**Fig. 4A-D**, and our previous reports (Mantegazza *et al*., 2017). In line with this observation, WT BMDCs treated with torin showed ASC speck formation away from the nucleus, similar to SLC15A4*^feeble^* DCs (**Fig. EV4C**). Altogether, our data indicate that mTORC1 signaling ensures optimal IL-1β secretion by limiting autophagy and promoting the assembly of the NLRC4 inflammasome at the perinuclear region where it appears excluded from autophagic membranes.

## Discussion

The molecular mechanisms underlying the regulation of inflammasome activity by phagosomal signaling are not completely elucidated (Moretti & Blander, 2014), nor are the pathways that bring together the processes of phagocytosis, autophagy and the downmodulation of inflammasomes. Autophagy has been increasingly recognized as an anti-inflammatory process (Deretic & Levine, 2018; Shi *et al*., 2012; Zhong *et al*., 2016). However, the molecular players that trigger autophagy after phagocytosis are only beginning to be unraveled. PRR stimulation by PAMPs/DAMPs on phagosomes is one mechanism proposed to trigger lysosomal and autophagy pathways by promoting expression of the coordinated lysosomal expression and regulation gene network (Adamson *et al*, 2016; Pastore *et al*, 2016). However, PRR stimulation also promotes inflammasome priming (Latz *et al*., 2013), leading to a pro-inflammatory response. Therefore, pro- and anti-inflammatory forces triggered after phagocytosis must be finely tuned according to the nature of the phagocytic cargo.

Here we show that the lysosomal histidine/peptide transporter SLC15A4 is recruited to phagosomes and phagosomal tubules in dendritic cells, as also observed for some PRRs (Blander, 2007; Mantegazza *et al*., 2012). Such an extensive recruitment surface may be required to ensure proper ligand or nutrient sensing and consequent cytosolic NOD2 signaling as observed in lysosomes (Nakamura *et al*., 2014). The role played by SLC15A4 in promoting type I interferon production and TLR9 signaling in pDCs and B cells has been extensively studied and supports the importance of this transporter in the pathogenesis of inflammatory disorders, including IBD (Baccala *et al*., 2013; Blasius *et al*., 2010; Kobayashi *et al*., 2021a; Kobayashi *et al*, 2021b; Kobayashi *et al*., 2014; Kobayashi *et al*., 2017; Sasawatari *et al*., 2011). However, the contribution of SLC15A4 to conventional DC function has been less studied (Nakamura *et al*., 2014). We now describe a previously unappreciated mechanism explaining the pro-inflammatory role of SLC15A4 in an *in vivo* model of DSS-induced colitis. Even though we do not rule out the contribution of other cell types to colon inflammation, we show that SLC15A4 promotes inflammasome activity and supports IL-1β production in intestinal conventional DCs by restraining autophagy, a process that genome-wide association studies have correlated with IBD (Larabi *et al*, 2020).

We also demonstrate that SLC15A4 promotes inflammasome activity triggered by an infectious stimulus, STm. The absence of SLC15A4 leads to decreased caspase-1 and GSDMD cleavage and decreased IL-1β production, which correlates with increased autophagy. Interestingly, decreased inflammasome activity triggered both by sterile and infectious stimuli in the absence of SLC15A4 was restored by the expression of constitutively active Rag B. Conversely, inhibiting mTORC1 decreased inflammasome activity in WT cells. Recent studies show that the Ragulator-Rag-mTORC1 axis is required for GSDMD oligomerization – post-cleavage – via ROS production (Evavold *et al*, 2021). Of note, these studies focus on post-cleavage events. Our observations, on the other hand, unravel a novel role for Rag B and mTORC1 in the regulation of inflammasome activity in the initial steps of inflammasome activation.

Inflammasome activity is also dependent on the site of inflammasome assembly (Magupalli *et al*, 2020; Martin *et al*, 2014). We previously observed that NLRC4 inflammasome formation at the perinuclear region appears to protect specks from sequestration by autophagic membranes (Mantegazza *et al*., 2017). We now show that the absence of SLC154 drives ASC speck formation away from the perinuclear region, and that remarkably, constitutively active mTORC1 rescues this aberrant phenotype. NLRP3 and pyrin inflammasome assembly occurs at the microtubule-organizing center (MTOC), and perinuclear positioning is dependent on histone deacetylase 6, an adaptor of the motor protein dynein (Magupalli *et al*., 2020). In contrast, NLRC4 inflammasome specks do not appear to localize at the MTOC and do not require microtubule transport for their activation ((Magupalli *et al*., 2020) and our unpublished observations). However, other cellular cytoskeletal filaments may be required for NLRC4 positioning. Similar to our observations in SLC15A4*^feeble^* BMDCs, peripheral ASC speck positioning is observed in the absence of the lysosomal adaptor AP-3 (Mantegazza *et al*., 2017). Interestingly, AP-3 – which recognizes a dileucine motif on SLC15A4 for lysosomal targeting – associates with the intermediate filament protein vimentin (Styers *et al*, 2005; Styers *et al*, 2004), which has been shown to interact with NLRP3 (dos Santos *et al*, 2015). Intermediate filaments also interact with motor proteins such as kinesins and dyneins (Helfand *et al*, 2004) and may be required to recruit these motors to sites of inflammasome assembly. Whether any of these associations are impaired in the absence of SLC15A4 and are relevant for NLRC4 assembly remains to be addressed. Furthermore, raptor, a component of mTORC1, is shown to associate with kinesins, dyneins and other molecules involved in cytoskeletal-filament assembly or function (Rabanal-Ruiz *et al*, 2021). Therefore, it is conceivable that reduced mTORC1 signaling caused by the absence of SLC15A4 impairs binding of ASC specks to cytoskeletal filaments or associated motor proteins that remain to be characterized. Regardless, our current observations support a novel and unexpected function for SLC15A4 and mTORC1 in ensuring NLRC4 inflammasome positioning at the perinuclear region, a location required for proper function and seemingly protected from autophagy, after phagocytosis (Mantegazza *et al*., 2017). We propose that the balance between inflammasome activation and deactivation by autophagy is regulated by recruitment of not only PRRs but also certain SLCs such as SLC15A4 to phagosomes. In turn, PAMPs/DAMPs and nutrient sensing performed by PRRs and SLCs link phagocytosis to mTORC1 signaling, the regulation of autophagy and the ultimate control of the immune response.

In homeostatic conditions, lysosomal mTORC1 signaling is turned on, keeping autophagy at bay. In the case of pathogenic microorganisms, in which the integrity of phagosomes is compromised, inflammasomes are triggered and autophagy is induced in response to PRR signaling, organelle damage or the pathogen itself (Deretic & Levine, 2018). In this scenario, SLC15A4 restrains autophagy to promote the anti-microbial response. Thus, SLC15A4 contributes to microbial sensing in phagosomes and allies with traditional PRRs to optimize immune responses against bacterial pathogens. In contrast, in the case of sterile inflammation such as IBD, SLC15A4 would play a detrimental role by sustaining inflammation and subsequent tissue damage. Whether this is a generalized mechanism and other phagosomal SLCs also contribute to the regulation of the phagosome-inflammasome-autophagy axis, certainly warrants further investigation.

## Materials and Methods

### Mice

C57BL/6 wild-type (WT) mice and SLC15A4*^feeble^* (C57BL/6J-*Slc15a4^m1Btlr^*) mice were originally purchased from The Jackson Laboratories (Bar Harbor, ME). SLC15A4*^feeble^* mice was previously described (https://mutagenetix.utsouthwestern.edu/phenotypic/phenotypic_rec.cfm?pk=426). Sex- and age-matched mice between 6 and 12 weeks of age were used in all experiments.

### Ethics Statement

Mice were bred under pathogen-free conditions in the Department of Veterinary Resources at the Children’s Hospital of Philadelphia or at Thomas Jefferson University, and were euthanized by carbon dioxide narcosis according to guidelines of the American Veterinary Medical Association Guidelines on Euthanasia. All animal studies were performed in compliance with the federal regulations set forth in the recommendations in the Public Health Service Policy on the Humane Care and Use of Laboratory Animals, the National Research Council’s Guide for the Care and Use of Laboratory Animals, the National Institutes of Health Office of Laboratory Animal Welfare, the American Veterinary Medical Association Guidelines on Euthanasia, and the guidelines of the Institutional Animal Care and Use Committees of Children’s Hospital of Philadelphia and Thomas Jefferson University. All protocols used in this study were approved by the Institutional Animal Care and Use Committee at the Children’s Hospital of Philadelphia (protocols #14-001064 and #16-001064) and Thomas Jefferson University (protocol #21-04-368).

### DSS-induced mouse model of colitis, colon sample preparation and analysis

WT and SLC15A4*^feeble^* male mice were randomized into control and experimental groups and co-housed at weaning across multiple cages. Experiments were performed three times with a total of 15 mice per genotype per experimental group and 12 male mice per genotype per control group. In experimental groups, 8 weeks-old mice were given 2.5% DSS (40,000–50,000 KDa molecular weight; Alfa Aesar J14489, Tewksbury, MA) in drinking water for 5 days, followed by normal drinking water for 10 more days. Age-matched control mice received water only. Body weight and stool appearance, consistency and presence of blood were recorded daily. Fecal occult blood was detected using Hemoccult single slides (Beckman Coulter Inc., Brea, CA). Blood stool index was determined using the following score: 0, normal feces, negative hemoccult; 1, soft but formed feces, positive hemoccult; 2, very soft feces, visible traces of stool blood; 3, diarrhea, rectal bleeding (Perse & Cerar, 2012; Wirtz *et al*, 2007). At day 16, mice were sacrificed, colon was dissected, length was measured, and Swiss rolls were prepared from PBS rinsed colon, fixed in 10% neutral buffered formalin (Polysciences) at room temperature for 48 h, rinsed in PBS and stored in 50% ethanol for histological staining and analysis. Haematoxilin and eosin staining, paraffin-embedding and histopathological analysis was performed at the Comparative Pathology Core, University of Pennsylvania, School of Veterinary Medicine. Histopathological analysis and scoring were performed blinded by expert veterinary pathologist Dr. Enrico Radaelli. Histopathological analysis was focused on assessing the mucosal changes (inflammatory and proliferative) of the mid and distal colorectal tract present in each sample (Gadaleta *et al*, 2017; Perse & Cerar, 2012). Briefly, scoring comprised the evaluation of mucosal/ crypt loss, crypt inflammation, mononuclear cells, neutrophils, epithelial hyperplasia and edema/ fibrosis.

### Reagents

LPS and monosodium urate crystals were purchased from InvivoGen (San Diego, CA) and alum was from ThermoFisher Scientific (Rockford, IL). TxR-conjugated OVA and EBSS were from Invitrogen (ThermoFisher Scientific). Mouse monoclonal anti-caspase-1 p20 (Casper-1) and rabbit polyclonal anti-ASC (AL177) were from Adipogen (San Diego, CA); rabbit polyclonal anti-LC3B (ab48394) and mouse monoclonal anti-p62 (ab56416) were from Abcam (Cambridge, MA); mouse monoclonal anti-β actin and mouse monoclonal anti-FLAG (clone M2) were from Sigma; rat monoclonal anti-CD40 (3/23), anti I-A^b^ (AF-120.1), anti-CD11c (HL3), anti-CD11b (M1/70), anti-CD86 (GL1) and anti-CD103 (M290) were from BD Biosciences (San Jose, CA); mouse monoclonal anti-GFP was from Roche (Indianapolis, IN); rabbit anti-mTOR (7C10), anti-phospho-mTOR (Ser2481), anti-p70 S6 kinase (49D7), anti-phospho-p70 S6 kinase (Thr389) and anti-phospho-S6 ribosomal protein (Ser235/236), and mouse anti-S6 ribosomal protein (54D2) were from Cell Signaling Technology (Danvers, MA). ELISA Ready-SET-Go! kits for mouse interleukin-6, was from eBioscience, (San Diego, CA); the anti-mouse-IL-1β ELISA set was from R&D (Minneapolis, MN). All secondary antibodies were from Jackson Immunoresearch (West Grove, PA). All restriction enzymes were from New England Biolabs (Ipswich, MA). Polymerase chain reaction (PCR) was performed using GoTaq kit (Promega, Madison, WI).

### Cell culture

Bone marrow cells were isolated and cultured for 7-9 d in RPMI-1640 medium (Gibco, ThermoFisher Scientific, Waltham, MA) supplemented with 10% low endotoxin-FBS (Hyclone, Logan, UT), 2 mM L-Gln, 50 μM 2-mercaptoethanol (Invitrogen) and 30% granulocyte-macrophage colony stimulating factor (GM-CSF)-containing conditioned medium from J558L cells (kindly provided by R. Steinman former laboratory, Rockefeller University, NY) for differentiation to DCs as described (Mantegazza & Marks, 2015; Winzler *et al*, 1997). DC2.4 mouse dendritic cell line (Chen *et al*, 2020; Shen *et al*, 1997) (kindly provided by Dr. Janis Burkhardt, Children’s Hospital of Philadelphia) was cultured in RPMI-1640 medium (Gibco, ThermoFisher Scientific) supplemented with 10% low endotoxin-FBS (Hyclone), 2 mM L-Gln and 50 μM 2-mercaptoethanol (Invitrogen).

Intestinal lamina propria was isolated after dissecting colon and dissociating the intestinal epithelium in Ca^2+^/Mg^2+^-free HBSS (Invitrogen) supplemented with 5% FBS and 2 mM EDTA for 20 min at 37°. Lamina propria was then digested by treatment with 1.5 mg/ml type VII collagenase (Sigma) and 40 μg/ml DNase I (Sigma) for 15 min at 37° to obtain single cell suspensions (as shown in http://www.jove.com/video/4040/; (Geem *et al*, 2012). Colonic DCs were then purified with anti-CD11c (N418) microbeads (Miltenyi Biotec Inc., Auburn, CA) (Geem *et al*., 2012).

### DNA retroviral constructs, retroviral production and transduction of dendritic cells

pMSCV-ASC-GFP, pLZRS-mCherry-LC3B and MigR1 retroviral constructs were kindly provided by Teresa Fernandes-Alnemri (Thomas Jefferson University, Philadelphia, PA), Erika Holzbaur (University of Pennsylvania, Philadelphia, PA) and Warren Pear (University of Pennsylvania), respectively. GFP was amplified using the forward primer 5’-ATCT**CTCGAG**ATGGTGAGCAAG GGCGAG-3’ (XhoI restriction site in bold) and reverse primer 5’-ATCT**GAATTC**TTACTTGTAC AGCTCGTC (EcoRI restriction site in bold) and sucloned between XhoI and EcoRI restriction sites of the MigR1-NotI vector (previously described (Lopez-Haber *et al*., 2020)) resulting in MigR1-NotI-GFP. hSLC15A4 was then cloned between BglII and XhoI restriction sites of the MigR1-NotI-GFP vector by digesting pLenti-hSLC15A4 (RC215932L2, NM_145648, OriGene, Rockville, MD) with BamHI and XhoI restriction enzymes resulting in MigR1-hSLC15A4-GFP. Retrovirus was produced by transfection of packaging cell line PLAT-E (Morita *et al*, 2000) (a generous gift of Mitchell Weiss, St. Jude Children’s Research Hospital, Memphis, TN) using Lipofectamine 2000 (Invitrogen, ThermoFisher Scientific) and harvested from cell supernatants 2 d later. 3 x 10^6^ BM cells were seeded on 6-well non-tissue culture treated plates per well for transduction 2 d after isolation, and transduced by spinoculation with 3 ml of transfected Plat-E cell supernatant in the presence of 8 μg/ml polybrene and 20 mM HEPES for 2h at 37°C. Retrovirus-containing media were then replaced with DC culture media. Puromycin (2 μg/ml) or the corresponding selection antibiotic was added 3 d after infection, and cells were collected for experiments 3 d later.

### shRNAs, lentiviral production and transduction of dendritic cells

pLKO.1-puromycin derived lentiviral vectors (Stewart *et al*, 2003) for small hairpin RNAs (shRNAs) against SLC15A4 and non-target shRNAs were obtained from the High-throughput Screening Core of the University of Pennsylvania. SLC15A4 #1 sense sequence: CCACCTGCATTACTACTTCTT; SLC15A4 #2 sense sequence: CCTCATTGTGTCTGTGAAGT A; SLC15A4 #3 sense sequence: CCAGAGTGTCTTCATCACCAA. Non-target sense sequence: GCGCGATAGCGCTAATAATTT. pLJM1-FLAG-RagB^Q99L^ and pLVX-FLAG-metap2 (control) were kindly provided by Dr. Roberto Zoncu (University of California, Berkeley, CA).

Lentivirus was produced by co-transfection of 293T cells (obtained from American Type Culture Collection, Mannassas, VA) with packaging vectors pDM2.G and pSPAx2 using calcium phosphate precipitation (Marks *et al*, 1995) and harvested from cell supernatants 2 d later. 3 x 10^6^ BM cells were seeded on 6-well non-tissue culture treated plates per well for transduction two d after isolation and transduced by spinoculation with 3 ml of transfected 293T cell supernatant in the presence of 8 μg/ml polybrene and 20 mM HEPES for 2h at 37°C (Savina *et al*, 2009). Lentivirus-containing media were then replaced with DC culture media. Puromycin (2 μg/ml) was added 3 d after infection, and cells were collected for experiments 3 d later.

For BM cell transduction with both retroviral constructs and lentiviral shRNAs, cells were first transduced with the indicated retroviral constructs, washed and subsequently transduced with the indicated lentiviral shRNAs. Lentivirus-containing media were then replaced with DC culture media. Puromycin (2 μg/ml) was added 3 d after infection, and cells were collected for experiments 3 d later. Only lentiviral shRNAs are puromycin resistant.

### Transfection of DC2.4 mouse dendritic cell line

DC2.4 were transfected with pEGFP-N1-hSLC15A4 (Nakamura *et al*., 2014) (kindly provided by Drs. Ira Mellman and Gerry Strasser, Genentech, South San Francisco, CA) using Neon transfection system (ThermoFisher Scientific) according to manufacturer’s instructions. Briefly, 2 x 10^6^ cells were washed in Ca^2+^/ Mg^2+^ free PBS and resuspended in 100 μl resuspension buffer R. Cells were then pulsed once with 5 μg of plasmid during 5 ms and 1,680 voltage and incubated overnight at 37°C.

### Bacterial strains and infections

For *in vitro* infections, STm were grown overnight in streptomycin containing LB medium at 37°C with aeration, diluted into fresh LB containing 300 mM NaCl, and grown standing at 37°C for 3 h to induce flagellin expression (Wynosky-Dolfi *et al*, 2014) unless otherwise indicated. Bacteria were washed with prewarmed Dulbecco’s Modified Eagle Medium (DMEM, Gibco), added to cells at a MOI of 5:1, and spun onto the cells at 200 x g for 5 min. Cells were incubated at 37°C in a tissue culture incubator with 5% CO_2_. Gentamycin (100 μg/ml) was added 1 h after infection and cells were harvested or analyzed by immunofluorescence microscopy or live cell imaging at the indicated time points.

### Real-time quantitative PCR (RT qPCR)

RNA was isolated from shRNA-transduced BMDCs using RNeasy kit (Qiagen, Germantown, MD). 1 μg RNA was reverse transcribed to cDNA using TaqMan™ Reverse Transcription Kit (Invitrogen, ThermoFisher Scientific). qPCR was performed in a StepOnePlus RT PCR System (Applied Biosystems, ThermoFisher Scientific) using TaqMan™ Fast Advanced master mix and TaqMan™ FAM probes (Applied Biosystems) according to manufacturer’s instructions. TaqMan™ FAM probes specific for Slc15a4 and the housekeeping genes β2-microglobulin (β2-m) and glyceraldehyde-3-phosphate-dehydrogenase (Gapdh) were Mm00505709_m1 (Slc15a4), Mm00437762_m1 (β2-m) and Mm99999915_g1 (Gapdh). PCR product formation was continuously monitored using the StepOnePlus™ Real-Time PCR System. Relative levels of Slc15a4 mRNA were calculated with the ΔΔCt method (Caino *et al*, 2011), normalized to the average of housekeeping genes and represented as mRNA fold change compared to control shRNA-treated cells. qPCR reactions were performed in triplicate. Experiments were independently performed three times.

### Inflammasome activation and measurement of cytokine production

200,000 BMDCs were seeded in triplicate in RPMI medium on 96-well round bottom plates, primed where indicated with 20 ng/ml of soluble LPS for 2-3 h, and then incubated with the indicated inflammasome stimuli (200 μg/ml alum, 200 MSU crystals or STm at MOI of 5:1), for 1 to 4 h. (Gross, 2012). Cells were pelleted at 200 x g at the indicated time points, supernatants were collected to measure cytokines using commercial ELISA kits (Gross, 2012).

For cytokine detection after *in vivo* DSS-treatment, isolated colonic DCs were cultured overnight, supernatants were collected to measure IL-1β by ELISA and cell pellets were lysed with Laemmli sample buffer with 2-mercapto-ethanol for immunoblotting analysis of LC3.

### Immunoblotting

For LC3, GSDMD and caspase-1, BMDCs were infected with STm in suspension, harvested by centrifugation at 200 x g at the indicated time points, and lysed in Laemmli sample buffer with 2-mercaptoethanol. Samples were then fractionated by SDS-PAGE on 12% or 15% polyacrylamide gels, transferred to PVDF membranes (Immobilon-P, Millipore) and analyzed using horseradish peroxidase-conjugated secondary antibodies (Jackson ImmunoResearch), enhanced chemiluminescence (GE Healthcare, Pittsburgh, PA) and iBright imaging system (ThermoFisher Scientific) or FluorChem R imaging system (ProteinSimple, Biotechne, San Jose, CA). For mTOR, p70 S6 kinase and S6 detection, samples were first lysed in lysis buffer containing 1% Triton X-100, 10 mM sodium pyrophosphate, 10 mM sodium β-glycerophosphate, 4 mM EDTA, 40 mM HEPES and EDTA-free protease inhibitors (Roche) at pH 7.4 (Chung *et al*, 2019) and then resuspended in Laemmli sample buffer with 2-mercaptoethanol. Samples were then fractionated on 6% (mTOR) or 12% (p70S6 kinase and S6) polyacrylamide gels, transferred to PVDF membranes (Immobilon-P, Millipore) and analyzed using horseradish peroxidase-conjugated secondary antibodies (Jackson ImmunoResearch), enhanced chemiluminescence (GE Healthcare, Pittsburgh, PA) and FluorChem R imaging system. Densitometric analyses of band intensity was performed using NIH Image J software, normalizing to control protein levels (Gross, 2012).

### Immunofluorescence microscopy and flow cytometry

Non-transduced BMDCs or BMDCs expressing SLC15A4-GFP or ASC-GFP and/or mcherry-LC3 and/or RagB^Q99L^ and metap2 (control), were infected with STm for the indicated time points, fixed with 3% formaldehyde in PBS, permeabilized with Permwash (BD Biosciences, San Jose, CA), and labeled with primary antibodies and Alexa Fluor-conjugated secondary antibodies (Jackson ImmunoResearch). Cells were analyzed by fluorescence microscopy using a Leica DMI6000 B inverted microscope, a Hamamatsu ORCA-flash 4.0 camera and Leica Microsystems Application Suite X software at the Department of Pathology at Children’s Hospital of Philadelphia or a Nikon A1R laser scanning confocal microscope with spectral detectors and Nikon NIS-Elements acquisition software at the Sidney Kimmel Cancer Center (SKCC) Bioimaging facility at Thomas Jefferson University. Images were analyzed using ImageJ (National Institutes of Health). LC3 fluorescence surrounding ASC specks was measured in a radio of 1 μm and normalized per unit area using Image J. ASC speck distance from nuclei was quantified using Analyze/Measure tools as detailed in Image J tutorial (https://imagej.nih.gov/ij/docs/pdfs/ImageJ.pdf).

BMDC phenotype was characterized by flow cytometry using an LSR-II and FloJo software (BD Biosciences). To assess ASC speck formation by flow cytometry, BMDCs were infected with STm for the indicated time points, fixed with 2% PFA in PBS for 10 m. Cells were finally washed in PBS with 0.5% BSA and 2 mM EDTA and analyzed by flow cytometry using a CytoFLEX (Beckman Coulter, Indianapolis, IN) and FlowJo software (BD Biosciences). Selection of activated and non-activated cell gates was performed using GFP-width and GFP-height parameters based on the differential shape of the fluorescence pulse depending on the fluorophore distribution within the cell, as previously described (Hoss *et al*., 2018; Sester *et al*., 2015) and shown in **Fig. EV3**.

### Live cell imaging

BMDCs or DC2.4 cells expressing SLC15A4-GFP were seeded on poly-L-lysine–coated glass-bottom 35-mm culture dishes (MatTek, Ashland, MA) on day 6 of culture. On day 7, cells were pulsed for 15 min with TxR-conjugated OVA (Invitrogen, ThermoFisher scientific) and LPS (100 μg/ml) coupled to 3-μm amino polystyrene beads (Polysciences Inc., Warrington, PA) as described previously (Mantegazza & Marks, 2015; Savina *et al*, 2010). DCs were then washed with RPMI, chased for 0– 2.5 h, and visualized using a Nikon A1R laser scanning confocal microscope with spectral detectors and equipped with a Tokai-Hit temperature and CO_2_-controlled chamber at the SKCC Bioimaging facility at Thomas Jefferson University. Images or videos were obtained with Nikon NIS-Elements acquisition software and analyzed using ImageJ (National Institutes of Health).

### Statistical analysis

Statistical analyses and data plots were performed using Microsoft Excel (Redmond, WA) and GraphPad Prism software (San Diego, CA). Statistical significance for *in vitro* experimental samples relative to untreated or control cells (as indicated) was determined using the unpaired Student’s t test and ANOVA after normality assessment using GraphPad. Statistical significance for mouse analyses and mouse samples in DSS-treated SLC15A4*^feeble^* mice relative to WT mice was determined using the non-parametric Mann-Whitney test (Fay & Proschan, 2010).

## Acknowledgments

We thank Julie Brill and Ira Mellman for critical reading of the manuscript; Roberto Zoncu, Erika Holzbaur, Cory Rogers, Teresa Fernandes-Alnemri and Emad Alnemri, Warren Pear, Ira Mellman, Gerry Strasser and Janis Burkhardt for the generous gifts of reagents; Michael S. Marks, the Department of Pathology at Children’s Hospital of Philadelphia and the Department of Microbiology and Immunology at Thomas Jefferson University for reagents, equipment and helpful discussions; María Yolanda Covarrubias at the Sidney Kimmel Cancer Center (SKCC) Bioimaging facility at Thomas Jefferson University and David Schultz and the High-throughput Screening core at the University of Pennsylvania for expert technical assistance, and the Flow Cytometry cores and Offices of Animal Resources at the Children’s Hospital of Philadelphia and Thomas Jefferson University.

This work was supported by NIH Grant R01 AI137173 (to C.L.-H. and A.R.M.) and R01 DK124369 (to K.E.H). The veterinary pathologists of the Penn Vet Comparative Pathology Core are partially supported by the Abramson Cancer Center Support Grant (P30 CA016520). The Aperio Versa 200 scanner used for whole slide imaging and the image analysis software was supported by an NIH Shared Instrumentation Grant (S10 OD023465-01A1). SKCC flow cytometry core is supported by NCI Core grant (P30 CA056036).

## Author contributions

Conceptualization: ARM

Methodology: CLH, XM, KEH, ARM

Investigation: CLH, ZH, ARM

Visualization: CLH, ZH, ARM

Funding acquisition: KH, ARM

Project administration: ARM

Supervision: KEH, ARM

Writing – original draft: ARM

Writing – review & editing: CLH, ZH, XM, KEH, ARM

---

**Figure EV1.**
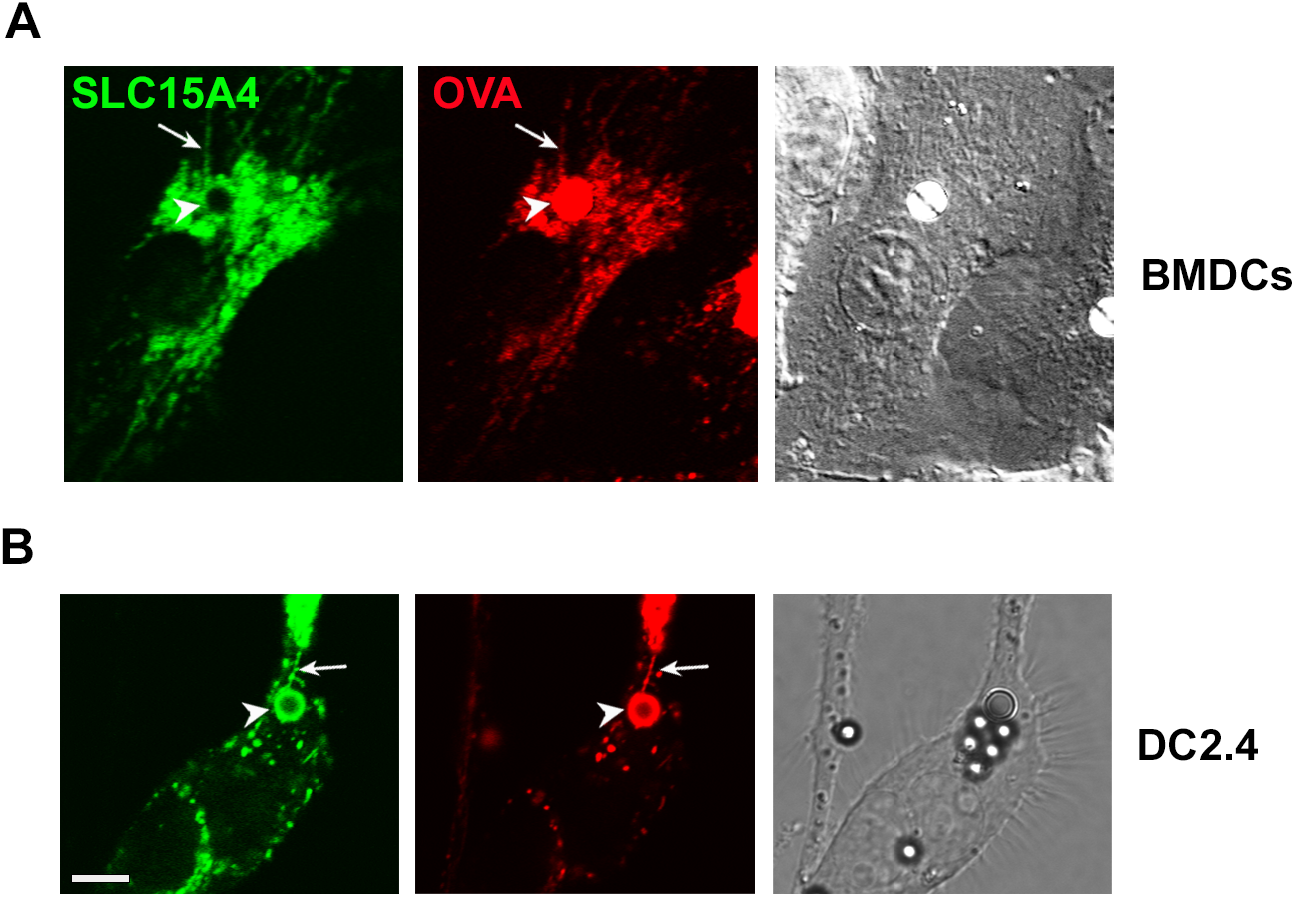
SLC15A4 is recruited to phagosomes and phagosomal tubules in DCs. WT BMDCs (**A**) or DC line DC2.4 (**B**) expressing SLC15A4-GFP were pulsed with LPS/OVA-TxR beads and analyzed by live cell imaging 2 h after the pulse. Representative images. DIC images show cell shape and outline. Arrowheads, phagosomes;

**Figure EV2.**
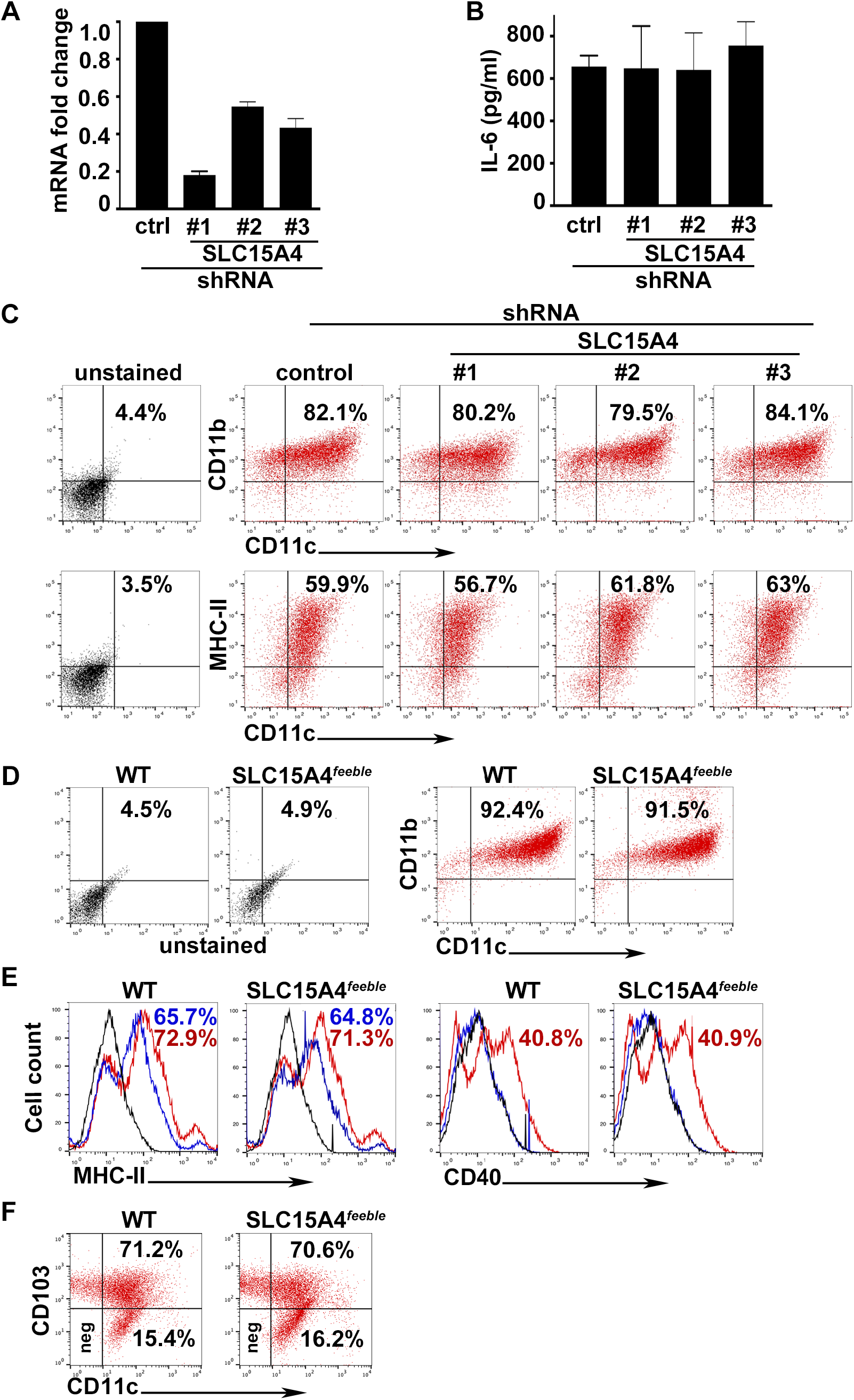
SLC15A4 knock-down or *feeble* mutation do not significantly impair DC differentiation, maturation or LPS priming. WT BMDCs transduced with lentiviruses encoding non-target (ctrl), or any of three SLC15A4 shRNAs (**A-C**), were untreated (**A, C**) or treated with soluble LPS for 3 h (**B**), or WT and SLC15A4*^feeble^* BMDCs were untreated (**D**) or treated with soluble LPS for 18 h (**E**), or intestinal DCs isolated from colon of WT or SLC15A4*^feeble^* mice were untreated (**F**). **A**. cDNA generated from isolated RNA was analyzed by RT-qPCR. Data from three independent experiments were normalized to the average of two housekeeping genes, and the ΔΔCt values were calculated and represented as mean ± SD fold change of mRNA in SLC15A4 shRNA-transduced cells relative to non-target ctrl-treated cells. **B**. Cell supernatants collected 3 h after treatment were assayed for IL-6 by ELISA. Representative plot of 3 independent experiments. (**C, D)**. Representative dot plots with the percentages of CD11b**^+^**/CD11c**^+^** BMDCs indicated as markers of DC differentiation after 7 days. **E.** Representative histograms with the percentages of CD40**^+^** or MHC-II**^+^** BMDCs as markers of DC maturation at day 7. Blue solid lines, untreated DCs; red solid lines, LPS-treated DCs; black solid lines, unstained controls. **F**. Representative dot plots with the percentages of CD11c**^+^**/CD103**^+^** DCs indicated as markers of intestinal DCs. arrows, phagosomal tubules. Scale bar, 6 μm.

**Figure EV3.**
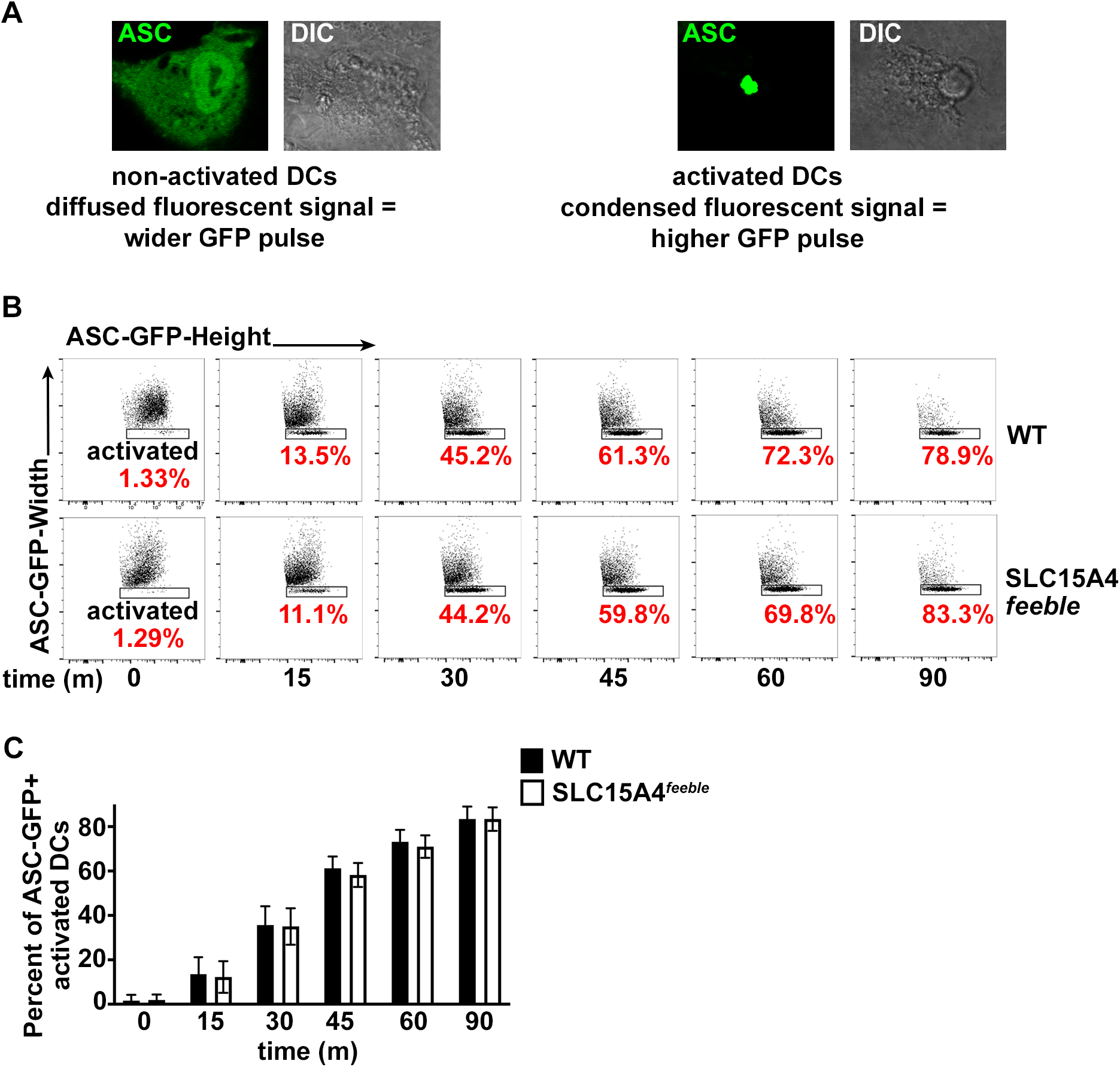
SLC15A4 does not affect the kinetics of ASC speck formation. WT or SLC15A4*^feeble^* BMDCs expressing ASC-GFP were infected with flagellin-expressing STm. Cells were fixed at the indicated time points after infection and analyzed by flow cytometry measuring GFP-width and GFP-height pulses. **A**. Cytosolic diffused ASC-GFP in non-activated cells correlate to a wider GFP signal compared to condensed ASC-GFP in activated cells. Representative images. DIC images show cell shape and outline. **B**. Representative dot plots, after gating on GFP^+^ cells. Note that the height of the GFP pulse is more prominent (*rectangular shape*) relative to the width of the GFP signal as the number of cells bearing ASC specks (activated DCs) increases overtime. Percent of GFP^+^ activated DCs are indicated in red. **C**. Plot represents three independent experiments. Data represent mean ± SD.

**Figure EV4.**
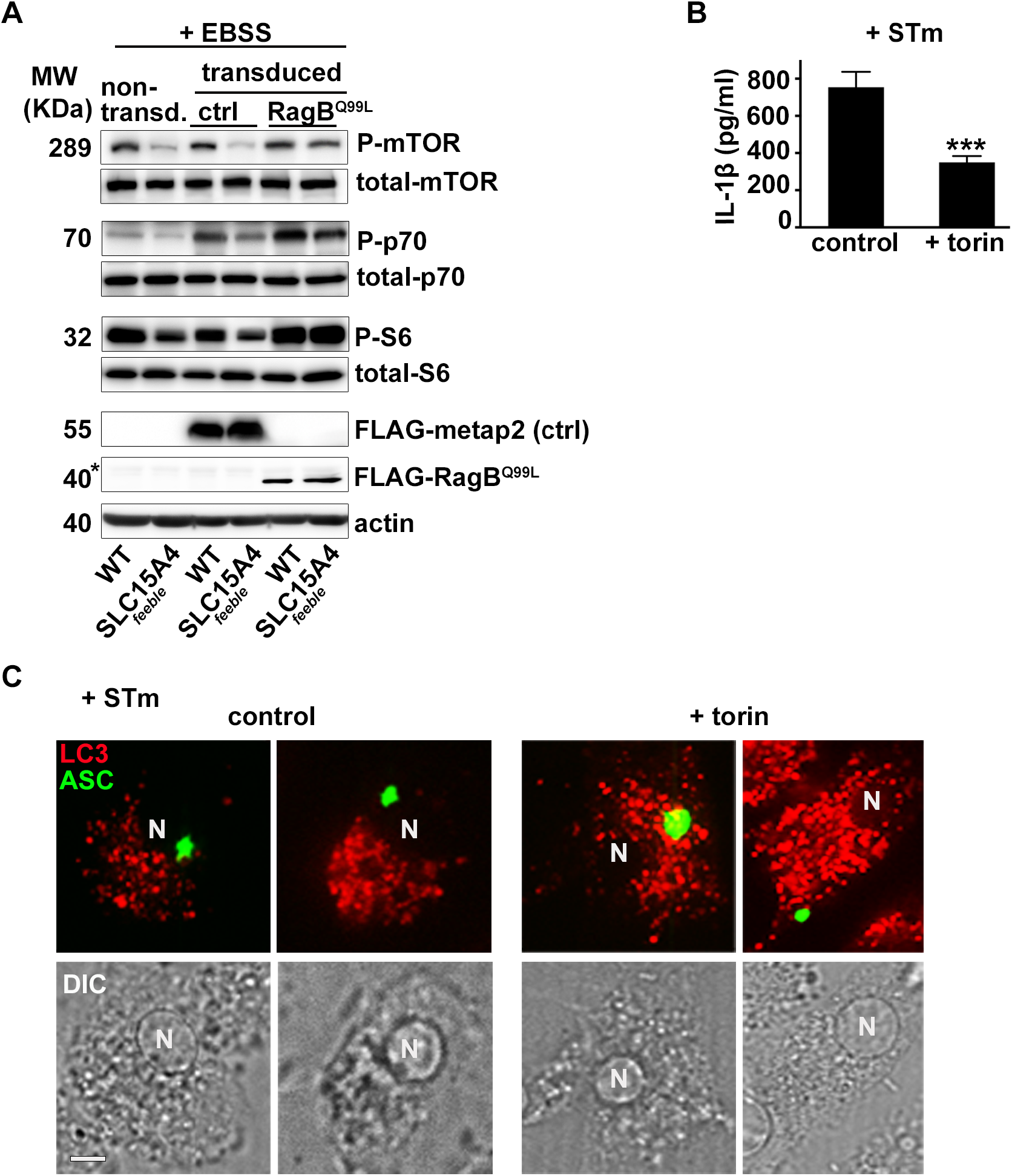
Constitutively active mTOR rescues defective mTOR signaling, and mTOR inhibition in WT DCs phenocopies SLC15A4*^feeble^* defects in inflammasome activity and positioning. WT or SLC15A4*^feeble^* BMDCs were untreated (**A-C**) or pre-treated with torin (**B**, **C**), transduced with FLAG-metap2 (ctrl) or FLAG-RagB^Q99L^ (A) or ASC-GFP and mcherry-LC3 (**C**) and starved in EBSS for 4 h (**A**) or infected with STm (**B**, **C**). **A**. Cell pellets were lysed, fractionated by SDS-PAGE and immunoblotted for phospho (P) and total mTOR, p70 kinase and S6, FLAG and actin. Representative immunoblots. Non-specific band right above FLAG-RagB^Q99L^ is indicated with an asterisk. **B**. Cell supernatants collected 1 h after infection were assayed for IL-1β by ELISA. Representative plot of 3 independent experiments. **C**. Cells were fixed and analyzed by fluorescence microscopy 1 h after infection. Representative images showing ASC speck (green) relative to LC3 (red). Note peripheral ASC speck positioning in torin-treated DCs. Corresponding DIC images show nuclear position. N, nucleus. Scale bar, 8 μm. Data represent mean ± SD. ***p<0.01.

## Expanded view Movie legends

**Movie EV1. SLC15A4 is recruited to phagosomes in BMDCs.** This movie shows the localization of SLC15A4-GFP (*green*) to phagosomal membranes and tubules after pulsing cells with LPS/OVA-TxR beads (*not colored*) and chasing for 2 h. Compression: jpeg. Frame rate: 7 fps.

**Movie EV2**. **SLC15A4 is recruited to phagosomes in DC2.4 cells.** This movie shows the localization of SLC15A4-GFP (*green*) to phagosomal membranes and tubules after pulsing cells with LPS/OVA-TxR beads (*not colored*) and chasing for 2 h. Compression: jpeg. Frame rate: 7 fps.

